# Exclusion of fire and grazing does not reduce annual grass invasion in a sagebrush steppe ecosystem

**DOI:** 10.64898/2026.05.21.726973

**Authors:** Joseph T. Smith, Brady W. Allred, Chad S. Boyd, Kirk W. Davies, Scott L. Morford, David E. Naugle, Thomas J. Rodhouse, Devin S. Stucki

## Abstract

Disturbance is widely recognized as a catalyst of invasion, but growing evidence suggests even protected communities are susceptible to severe infestation. We used kīpukas—naturally isolated patches of minimally-disturbed vegetation surrounded by lava flows—as a large-scale natural experiment to test the long-term biotic resistance of protected sagebrush ecosystems threatened by *Bromus tectorum* and other invasive annual grasses. Employing a robust causal inference approach combining matching with regression adjustment, we compared protected communities within kīpukas to otherwise similar communities exposed to contemporary disturbance regimes. Despite their near-total protection from fire and livestock grazing, kīpukas were extensively invaded by annual grasses (18.9 ± 0.28% cover), with abundance comparable to or slightly exceeding disturbed sites (16.7 ± 0.29% cover). These findings challenge the notion that protection from disturbance confers effective long-term resistance to invasion, instead demonstrating that invaders can establish, proliferate, and drive ecosystem transformation where favorable abiotic conditions prevail. Our findings reveal the limits of passive protection as a conservation strategy, suggesting active management may be necessary to prevent ecosystem degradation by aggressive invaders.

## Introduction

Disturbance is believed to play a central role in the invasibility of ecosystems (Elton 1958, Hobbs and Huenneke 1992). By damaging or destroying resident individuals, disturbance frees up space (Tilman 1994) and resources (Davis et al. 2000), facilitating establishment and growth of new individuals. Non-native species often benefit disproportionately from disturbance (Jauni et al. 2015) due to enemy release (Hierro et al. 2006), novel weapons (Callaway and Ridenour 2004), or the overrepresentation of ruderal species among exotic species pools (Catford et al. 2012, Pearson et al. 2023). The close link between invasion and disturbance has led some to propose that invasive species may be incidental “passengers” of external stressors such as human land use and altered disturbance regimes, rather than “drivers” of ecosystem degradation (MacDougall and Turkington 2005, Hille Ris Lambers et al. 2010).

Yet, invasive species can pose significant challenges even among minimally-disturbed ecosystems. Invasive plants, in particular, are a widespread and growing problem among protected areas globally (Shackleton et al. 2020, Carneiro et al. 2024), in some cases altering ecosystem function in profound ways (Vitousek et al. 1987, Foxcroft et al. 2017, Ren et al. 2021). Although disturbance may explain some invasions, other invaders are clearly capable of driving change.

The transformation of western North American shrublands by invasive annual grasses is a phenomenon closely associated with anthropogenic disturbance. Cheatgrass (*Bromus tectorum* L.), an annual grass introduced from Eurasia in the late 19th century, now dominates millions of hectares in the Intermountain region where it fuels larger and more frequent wildfires (D’Antonio and Vitousek 1992, Fusco et al. 2019), degrades habitat for sensitive species (Coates et al. 2016, Hobart et al. 2026), and alters ecosystem processes such as water and nutrient cycling (Sperry et al. 2006, Hooker et al. 2008, Schaeffer et al. 2012). Failing to fit neatly within either the passenger or driver category, cheatgrass has been described as a “back-seat driver” of ecosystem change (Bauer 2012). This label refers to its alleged reliance on disturbance for establishment, and its ability to subsequently drive change via a positive feedback with fire. Disturbance of plants or soil is thought to provide cheatgrass an initial foothold within sagebrush communities (Leffler et al. 2016), and the extensive degradation of the native perennial bunchgrasses through unregulated livestock grazing and deliberate burning in the late 19th and early 20th centuries is widely credited with enabling its explosive initial spread (Leopold 1941, Young et al. 1972, Mack 1981). Observational studies of contemporary patterns in cheatgrass abundance have emphasized continued impacts of livestock grazing and fire as drivers of invasibility (Reisner et al. 2013, Condon and Pyke 2018, Mahood and Balch 2019, Root et al. 2020, Williamson et al. 2020).

Although disturbance can undoubtedly accelerate invasion, recent research suggests its role in driving ecosystem transformation by cheatgrass has been overstated. Smith et al. (2023) used remote sensing-derived fractional vegetation cover data to show that transitions to annual grass dominance in the Great Basin over the last three decades more often preceded than followed fire. Similarly, Urza et al. (2024) combined extensive field observations with historical fire records to demonstrate that abiotic suitability drives overall risk of annual grass dominance despite fire acting as a catalyst. These large-scale studies lend considerable weight to decades of anecdotal observations of cheatgrass established within long-undisturbed plant communities (Daubenmire 1940, Passey and Hugie 1963, Tisdale et al. 1965, Bangert and Huntly 2010). Although often noted as a minor component of undisturbed communities, severe and persistent infestations of cheatgrass have also been documented at protected sites (Driscoll 1964, Passey et al. 1982, Kindschy 1994, Tausch et al. 1994, Gale and Goodrich 1999).

Although these studies suggest cheatgrass can invade and substantially alter plant communities even in the absence of disturbance, key uncertainties remain. Could disturbances other than fire explain the extensive increases in cheatgrass dominance across the west? Livestock grazing, though now regulated, remains a ubiquitous land use across the sagebrush biome and may represent a significant departure from the disturbance regimes under which many sagebrush-bunchgrass communities evolved (Mack and Thompson 1982). Experimental studies of modern grazing practices have generally found similar or greater invasion when livestock are excluded (Davies et al. 2009, Porensky et al. 2020, Copeland et al. 2023). However, even sites protected from grazing today—e.g., in protected areas or exclosures—may express persistent legacies of past disturbance, obscuring the extent to which these patterns reflect the inherent vulnerability of these plant communities.

Resolving this uncertainty is central to understanding ecosystem invasibility and to the effective conservation of remaining sagebrush ecosystems. Cheatgrass is one of North America’s most consequential invaders, responsible for substantial ecological and economic losses (Germino et al. 2016). Its expansion, and that of other invasive annual grasses, is now regarded as the principal threat to sagebrush ecosystems and their unique biota (Doherty et al. 2022). If cheatgrass is capable of driving ecosystem transformation without disturbance, then conservation strategies relying on passive protection are unlikely to succeed and proactive intervention may be needed to prevent continued degradation.

To address the extent to which protection from disturbance confers lasting resistance to cheatgrass invasion, we took advantage of the unique natural isolation of kīpukas—relict patches of vegetation surrounded by lava flows. Hundreds of kīpukas containing sagebrush plant communities occur among lava flows in the northern Great Basin floristic province (Fig. 1), a region long impacted by cheatgrass and other invasive annual grasses. Owing to the rugged and mostly unvegetated surrounding lava, many of these kīpukas escaped historical overgrazing and are largely protected from wildfire (Passey and Hugie 1963), thus presenting a replicated natural experiment to assess the cumulative effects of these disturbances on contemporary invasion severity. We applied matching methods (Ho et al. 2007, Stuart 2010) to select a set of protected sites within kīpukas and a comparable set of nearby sites exposed to contemporary disturbance regimes (i.e., livestock grazing and frequent wildfire). We then quantified severity of invasion at all sampled sites using remote sensing-derived estimates of fractional vegetation cover and estimated effects of protection within a potential outcomes causal framework (Rubin 1974, Splawa-Neyman et al. 1990).

**Figure 1:**
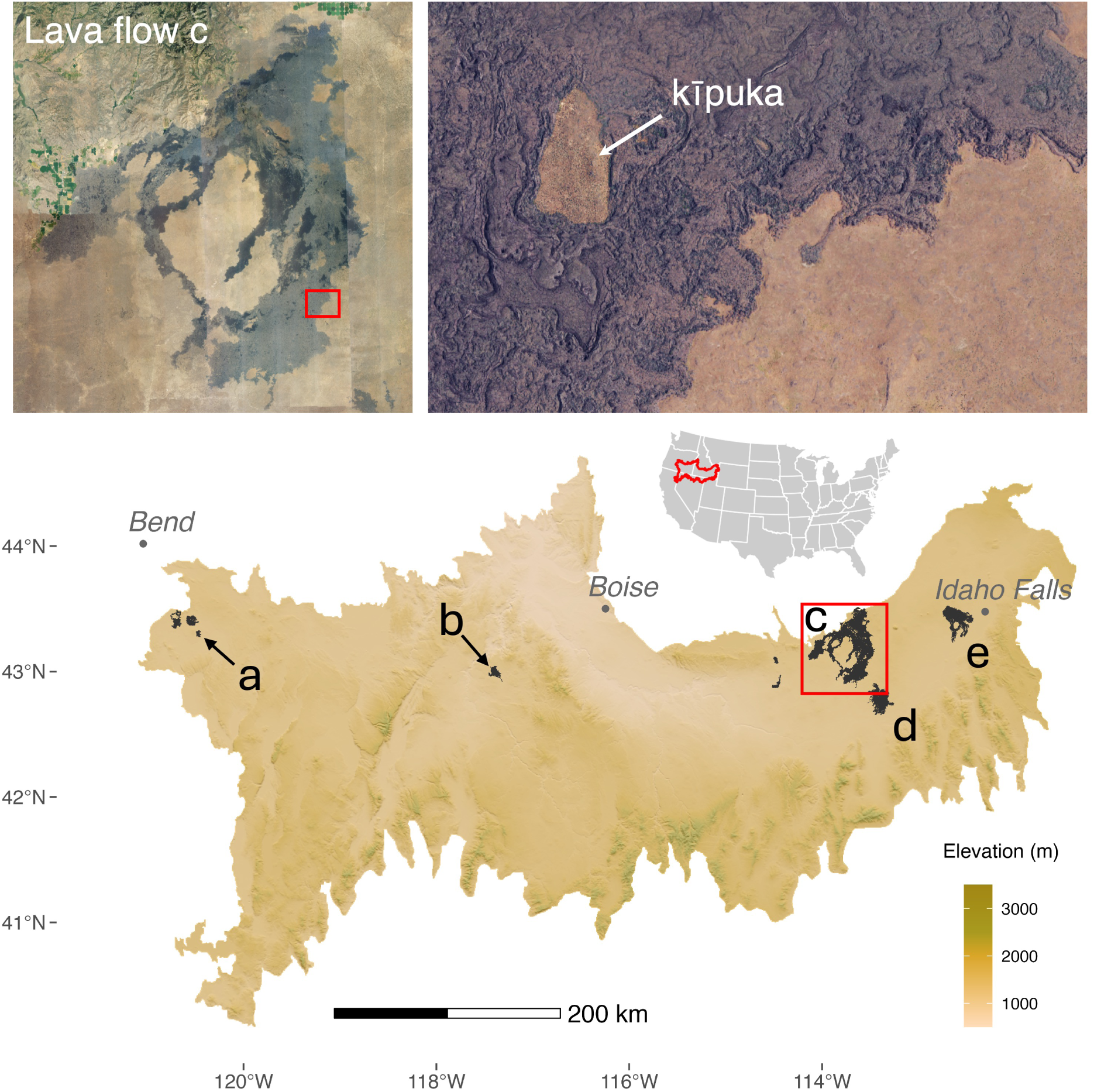
Study area in the northern Great Basin, USA. Lava flows (black polygons labeled a–e) containing kīpukas occurred along a narrow latitudinal band; elevations ranged from 1,299 to 1,772 m. Top left inset depicts lava flow *c*, the larger of two lava flows in Craters of the Moon National Monument and Preserve, Idaho. The top right inset depicts a kīpuka on the eastern border of lava flow *c*

## Methods

### Study area

We searched for kīpukas among lava flows in the Northern Basin and Range and Snake River Plain ecoregions (Level III; Omernik and Griffith 2014). We obtained a database of digitized kīpukas within Craters of the Moon National Monument and Preserve (CRMO; c and d in Fig. 1) from the National Park Service. Additionally, six large lava flows outside CRMO were systematically searched for kīpukas with the aid of aerial imagery. All potential kīpukas were individually inspected using aerial imagery from Google and the National Aerial Imagery Program (NAIP; 0.5 or 0.6 m resolution) to identify and discard candidates that were incompletely isolated by lava, in close proximity (<200 m) to the outer perimeter of the lava flow, intruded by roads or trails, or otherwise misidentified in the initial search.

The boundaries of all screened kīpukas were digitized in Google Earth Engine.

### Analysis mask

To focus our analysis on natural vegetation communities and avoid confounding factors, we developed a mask to eliminate open water, developed areas, lava, cropland, forests and woodlands, roads, and recently burned areas (i.e., burned during the period represented in the remotely sensed vegetation data) from sampling. We manually labeled 400 points with one of 8 landcover classes: shrubland/grassland, woodland/forest, cropland, water, lava flow, cinder cone, barren, and other (including roads and developed areas). We then used these labels to train a k-nearest neighbors classifier, using the AlphaEarth Foundations satellite embeddings (Brown et al. 2025) as predictors (year = 2020). Briefly, the satellite embeddings compress large quantities of spatial and temporal information from multiple Earth observation satellites into a set of 64 features or “embeddings,” that are conceptually similar to ordination axes. Embeddings are provided at an annual time-step and 10 m resolution, with a global extent. The resulting classification had an average cross-validation multi-class accuracy of 93.6 ±1.1% (κ = 0.916 ±0.015). Predicted classes other than shrubland/grassland were excluded from sampling. We further masked areas within 100 m of a road using the US Census Tiger/Line dataset and areas that burned during the period 2020–2024 using the Monitoring Trends in Burn Severity (MTBS) dataset (Eidenshink et al. 2007), which only maps fires >400 ha and the MODIS MCD64A1 Burned Area Product (500 m; NASA LP DAAC, 2025), which is capable of detecting fires as small as ∼100 ha.

### Sampling and matching

Kīpukas that passed visual screening were buffered inward by 15 m to avoid pixel overlap with the surrounding lava and sampled on a regular point grid. The sampling rate was based on the area of the kīpuka, with smaller kīpukas sampled at a higher density than larger kīpukas. Unprotected areas outside the lava flows but within 5 km of a sampled kīpuka were sampled on a regular 240 m point grid, ensuring similar soil development history as kīpukas primarily represent pre-Holocene surfaces isolated by relatively recent (∼2–15 ka) lava flows. We extracted a suite of topographic and edaphic covariates (Table 1) from sampling points that did not intersect the analysis mask; we used these covariates to match protected sites to an environmentally similar group of unprotected sites.

**Table 1:** Variables used to match protected samples to unprotected samples. The Mahalanobis distance was used to measure dissimilarity.

| Variable | Source | Resolution |
| --- | --- | --- |
| Elevation (m) | USGS 3DEP DEM | 10 m |
| Slope (degrees) | USGS 3DEP DEM | 10 m |
| Northness:<br>$\cos(\pi \times \text{aspect}/180)$ | USGS 3DEP DEM | 10 m |
| Eastness: $\sin(\pi \times \text{aspect}/180)$ | USGS 3DEP DEM | 10 m |
| Continuous heat-insolation<br>load index | Ecologically Relevant Geomorphology dataset<br>(Theobald et al. 2015) | 10 m |
| Multi-scale topographic<br>position index | Ecologically Relevant Geomorphology dataset<br>(Theobald et al. 2015) | 10 m |
| Lithology (parent material)<sup>†</sup> | Ecologically Relevant Geomorphology dataset<br>(Theobald et al. 2015) | 90 m |
| Depth to bedrock (cm)<sup>‡</sup> | Soil Landscapes of the United States<br>(SOLUS100) dataset (Nauman 2024) | 100 m |
| Clay fraction, 0-30 cm<sup>‡</sup> | Soil Landscapes of the United States<br>(SOLUS100) dataset (Nauman 2024) | 100 m |
| Sand fraction, 0-30 cm<sup>‡</sup> | Soil Landscapes of the United States<br>(SOLUS100) dataset (Nauman, 2024) | 100 m |
| Silt fraction, 0-30 cm<sup>‡</sup> | Soil Landscapes of the United States<br>(SOLUS100) dataset (Nauman, 2024) | 100 m |

<sup>†</sup> Exact match required.
<sup>‡</sup> Used only for matched Set 2 (see Methods for details).

The objective of matching was to produce an overall sample in which the protected and unprotected samples were balanced with respect to covariates likely to affect either the outcome (invasive annual grass cover) or the probability of “treatment” (i.e., being protected within a kīpuka). This balance removes systematic differences between the protected and unprotected samples that could otherwise confound the effect of the treatment, and is what enables causality to be ascribed to protection. The logic is the same as that underlying randomized experiments: covariate balance ensures that the potential outcomes are independent of the treatment assignment (Rubin 1974).

Because protected and unprotected samples were located nearby and were likely to experience similar weather, topography and soils were the primary covariates that we sought to balance. We required exact matching on lithology to ensure comparable parent material between protected and unprotected sites. Additional topographic variables selected for matching included elevation, slope, and aspect (expressed as northness and eastness), the continuous heat-insolation load index (Theobald et al. 2015); and the multi-scale topographic position index (Theobald et al. 2015). Edaphic variables included parent material (lithology; Theobald et al. 2015), basic attributes of soil texture (sand, silt, and clay fractions), and depth to bedrock (SOLUS100; Nauman et al. 2024).

Because SOLUS100 soil data (100 m resolution) were too coarse for small kīpukas (<3 ha), we prepared two matched sets. Set 1 included samples from all kīpukas regardless of size, but omitted continuous soil variables from the matching and regression adjustment steps. Set 2 used all available soil variables for matching and regression adjustment, but only included kīpukas ≥ 3 ha. Our interpretation focuses primarily on Set 1 because soil properties have relatively weak effects on invasive annual grass abundance in the northern Great Basin (Bansal and Sheley 2016), and we considered the greater replication achieved in Set 1 an asset for inference.

We selected matches without replacement using a greedy nearest neighbor algorithm (Ho et al. 2007, Stuart 2010), and used the Mahalanobis distance as a metric of dissimilarity. Briefly, this involved iterating through all of the protected samples one at a time, selecting the most similar unprotected sample as its match, and then removing that selected sample from the pool of potential matches. Matching was performed using the *MatchIt*package (Ho et al. 2011) in R. Because the algorithm is sensitive to the order of the data, we repeated the procedure 1,000 times, randomly shuffling the protected samples at each iteration before running the algorithm (Wauchope et al. 2022). We selected the match that produced the smallest maximum standardized difference in means (i.e., the difference between the means of the protected and unprotected samples divided by their pooled standard deviation) across covariates for analysis. We considered the matched dataset adequately balanced when standardized differences in means were less than 0.25 and variance ratios fell between 0.5 and 2 for all covariates (Rubin 2001, Stuart 2010).

### Outcome data

We used data from the Rangeland Analysis Platform (RAP; Allred et al. 2025) to derive fractional cover of key vegetation functional groups at the selected set of matched spatial samples. RAP’s 10 m, Sentinel-2-based fractional cover dataset provides annual (beginning in 2018), 10 m-resolution gridded estimates of cover of invasive annual grasses (IAG), perennial forbs and grasses (PFG), shrubs (SHR), and bare ground (BGR) among other functional groups. IAG primarily represents cheatgrass and medusahead (*Taeniatherum caput-medusae* (L.) Nevski) in our study area (see Appendix S1). Perennial bunchgrasses and shrubs are the co-dominant functional groups comprising intact sagebrush vegetation communities, and bare ground may play an important role in these communities by inhibiting the spread of fire (Brooks et al. 2001). We therefore extracted estimates of IAG, PFG, SHR, and BGR at each sample. We averaged over the 5 most recent years available in the RAP dataset (2020–2024) to reduce the influence of interannual fluctuations due to precipitation variability (Rinella et al. 2024). Remote sensing of fractional vegetation cover is a new and rapidly evolving field involving greater uncertainty than field-based methods; thus, we include a sensitivity analysis in Appendix S2 comparing the results obtained from three independent fractional cover datasets.

### Outcome model

For each plant functional type, we fit a regression model relating the outcome—percent cover—to the treatment (protected = 1, unprotected = 0) and the environmental covariates in Table 1. We fit linear mixed effects models with kīpukas nested within lava flows as random effects. Fixed effects included treatment, all covariates (z-standardized), and two-way interactions between the treatment and each of the covariates. Inference was not directly based on coefficient estimates and their standard errors; rather, the fitted model was used to estimate counterfactual outcomes for each protected sample, which were subsequently used to estimate the causal effect of protection (see *Causal effect estimation*, below). This combination of matching and regression adjustment is considered “doubly robust” to errors in matching and model misspecification (Ho et al. 2007), as matching reduces model dependence and regression adjustment compensates for residual imbalance in the matched dataset. We fit regression models with the **lmer** function in the *lme4* package (Bates et al. 2015) in R using the restricted maximum likelihood.

### Causal effect estimation

We estimated the average treatment effect among the treated (ATT), which is the difference between observed cover in protected kīpukas and their predicted cover had they been unprotected, using the *marginaleffects* package (Arel-Bundock et al. 2024) in R. Standard errors and confidence intervals were calculated using the nonparametric block bootstrap (n = 5,000 iterations) with matched pairs as resampling units (Abadie and Spiess 2022)

### Bias and error correction

Remote sensing-derived fractional cover estimates contain substantial prediction error, yet statistical inference based on these data can yield biased parameter estimates with artificially high precision due to the ease with which very large sample sizes are achieved. To account for this uncertainty, we applied the multiple imputation method of Proctor et al. (2023), which adjusts estimates and confidence intervals to reflect the relationship between remotely sensed and field measured cover. Using 334 monitoring plots sampled from 2018–2023 (details in Appendix S1), we compiled a regional calibration dataset with co-located remotely sensed (RAP) and ground-estimated (line point intercept) cover for the four response variables. We then repeatedly: (i) drew a bootstrap sample of the calibration dataset, (ii) fit a regression linking RAP estimates (independent variable) to ground-estimated cover (dependent variable), and (iii) used the fitted model and residual standard deviation to draw plausible cover estimates for all samples. This procedure was repeated 5,000 times concurrently with the block bootstrap (see *Causal effect estimation*), ensuring standard errors and confidence intervals incorporated uncertainty from both remote sensing error and the matching process.

## Results

We initially identified 484 kīpukas, of which 258 passed visual screening and met our inclusion criteria. Samples were obtained from 197 of these, ranging in size from 0.26–449 ha (median = 1.5 ha), from across 5 lava flows (Fig. 1). The remaining 61 kīpukas were not represented in the sample due to insufficient interior area (i.e., no suitable pixels intersected the inner buffer). Only 6.5% of kīpukas ≥200 m from the edge of a lava flow were excluded from sampling due to recorded fire since 1984, confirming the relative rarity of fire in these communities.

Sample Set 1, representing 197 kīpukas across 5 lava flows, included 1,351 protected samples and an equal number of matches selected from a pool of 23,390 unprotected samples. 71% of unprotected samples in Set 1 burned at least once, and 40% burned more than once, since 1984. Sample Set 2, representing the subset of kīpukas ≥3 ha in size (n = 79 from 4 lava flows), included 782 protected samples and their matched unprotected samples. 67% of unprotected samples in Set 2 burned at least once, and 38% burned more than once, since 1984. Kīpukas were represented by a median of 6 samples each (IQR = 3–9). Samples spanned a narrow latitudinal band between 42.67°N and 43.57°N and elevations from 1,299–1,772 m encompassing typical big sagebrush (*Artemisia tridentata* Nutt.) elevations in the Northern Basin and Range and Snake River Plain ecoregions (Fig. 1).

Set 1 was well-balanced, with all covariates achieving absolute standardized differences in means <0.11 and variance ratios between 0.99 and 1.33. Estimated cover of annual forbs and grasses, perennial forbs and grasses, bare ground, and shrubs among pairs of matched samples in Set 1 is shown in Figure 2. Set 2 was mostly well-balanced with the exception of depth to bedrock. The standardized difference in means for depth to bedrock after matching was -0.6 (i.e., 0.6 standard deviations, or roughly 15 cm, shallower among protected samples). This residual imbalance is not necessarily problematic for estimating the ATT, however, because depth to bedrock was also included as a predictor in the regression used to estimate potential outcomes.

**Figure 2:**
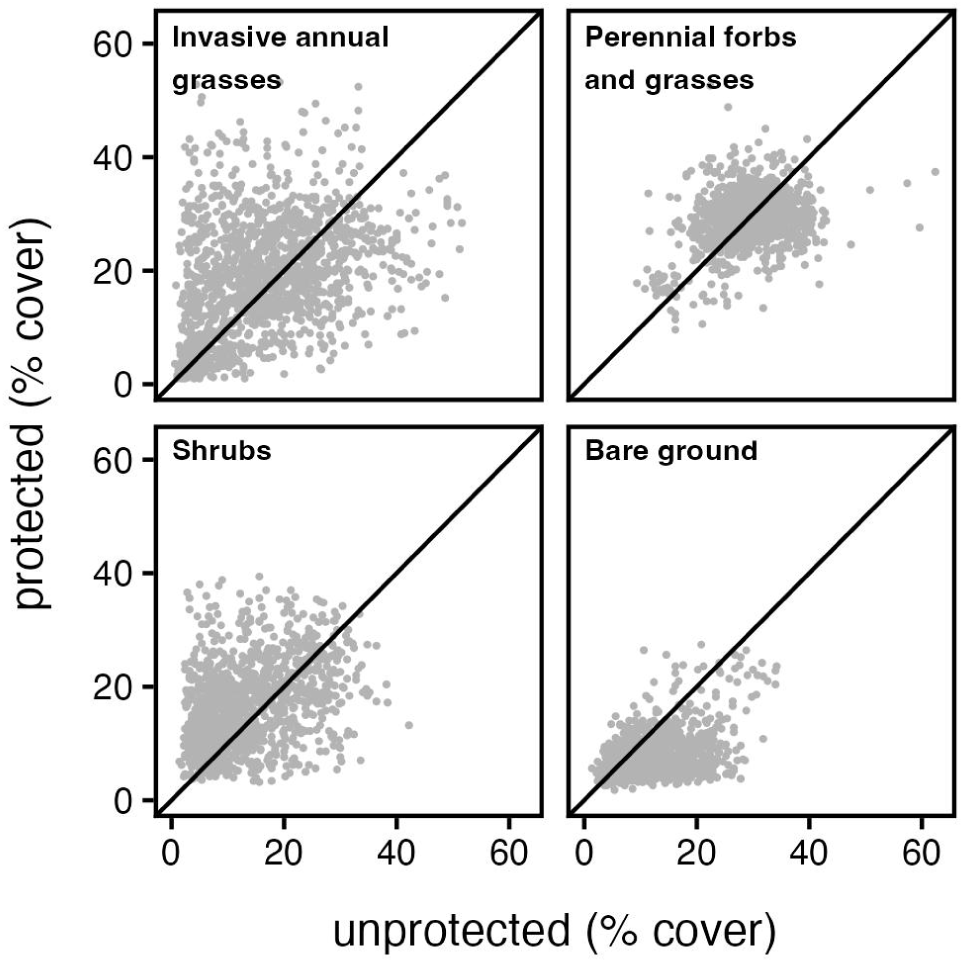
Remote sensing-based estimates of percent cover of four plant functional types among matched protected and unprotected samples (n = 1,351 matched pairs). Points above the diagonal indicate higher cover in protected samples; points below indicate higher cover in unprotected samples.

Cover of invasive annual grasses was 16.7 ±0.29% (mean ±SE) among unprotected samples and 18.9 ±0.28% among protected samples. Cover of perennial forbs and grasses was 29.2 ±0.15% among unprotected samples and 29.1 ±0.12% among protected samples. Cover of shrubs was 12.8 ±0.21% among unprotected samples and 16.2 ±0.19% among protected samples. Cover of bare ground was 12.0 ±0.16% among unprotected samples and 7.4 ±0.10% among protected samples.

The average effect of protection (95% confidence interval) was 1.51 (0.37 to 2.64)% for invasive annual grasses, 0.01 (−1.23 to 1.24)% for perennial forbs and grasses, -3.80 (−4.38 to -3.23)% for bare ground, and 2.80 (1.98 to 3.64)% for shrubs (Fig. 3). Using the subset of kīpukas with soils data (Set 2), these effects were 1.34 (−0.35 to 3.08)% for invasive annual grasses, -0.14 (−2.06 to 1.79)% for perennial forbs and grasses, -2.73 (−3.50 to -1.98)% for bare ground, and 2.68 (1.48 to 3.89)% for shrubs (Fig. 3).

**Figure 3:**
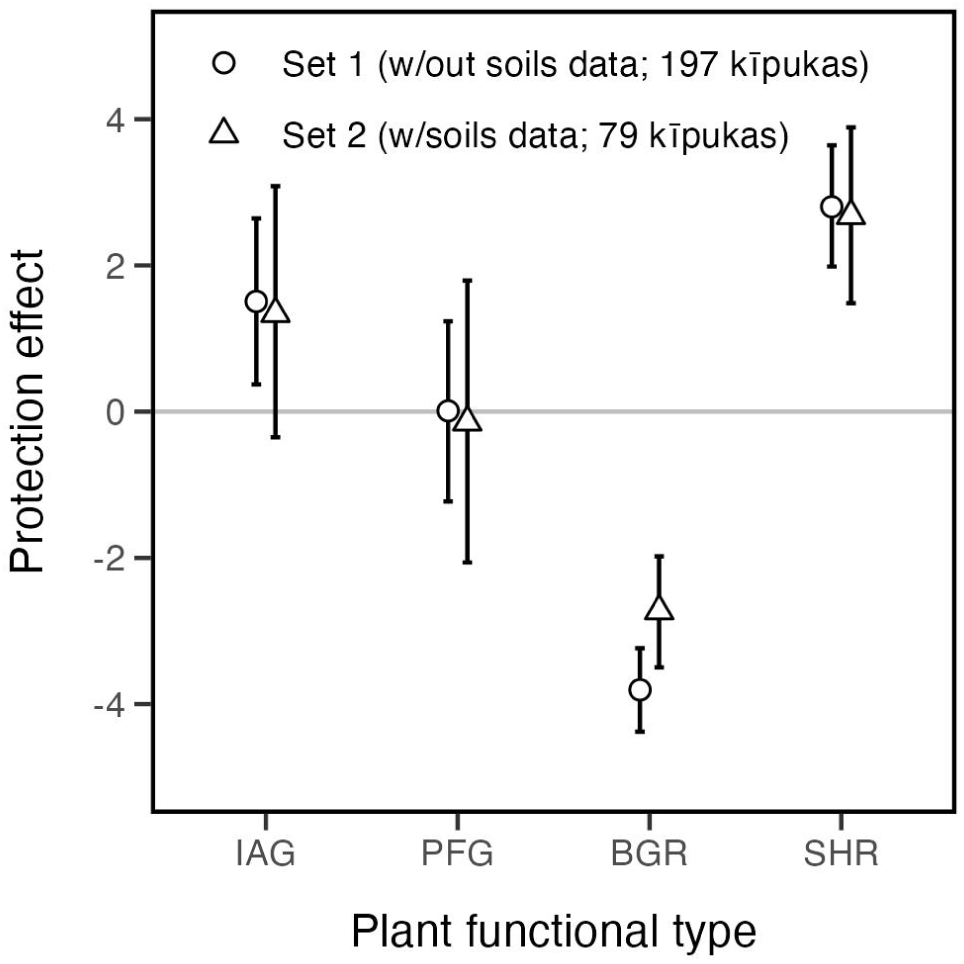
Effect of protection from disturbance (estimate and 95% confidence interval) on invasive annual grass cover (IAG), perennial forb and grass cover (PFG), bare ground cover (BGR), and shrub cover (SHR) in sagebrush plant communities of the northern Great Basin. Effect size is in units of percent cover. Positive (negative) values indicate protection from disturbance increased (decreased) cover relative to sites exposed to more typical contemporary disturbance regimes, including livestock grazing and elevated fire frequency.

## Discussion

This study is the first to use a quasi-experimental approach to quantify the effect of disturbance on the invasibility of sagebrush ecosystems by annual grasses. Isolated by lava flows thousands of years before the arrival of the invaders, sagebrush communities within kīpukas offer a rare opportunity to observe the invasibility of these ecosystems in a minimally-disturbed state. Contrary to the view that disturbance is a prerequisite for severe infestation, invasive annual grasses were widespread throughout these uniquely protected communities. In fact, protection appeared to increase rather than decrease invasion severity. In the full dataset, protection increased cover of annuals by about 1.5% on average. This effect was slightly attenuated and estimated with greater uncertainty among the smaller subset of kīpukas with soils data, but directionally unchanged. In either case, annual grasses are clearly capable of both establishing and proliferating in minimally-disturbed sagebrush plant communities where favorable abiotic conditions prevail (Fig. 4).

**Figure 4:**
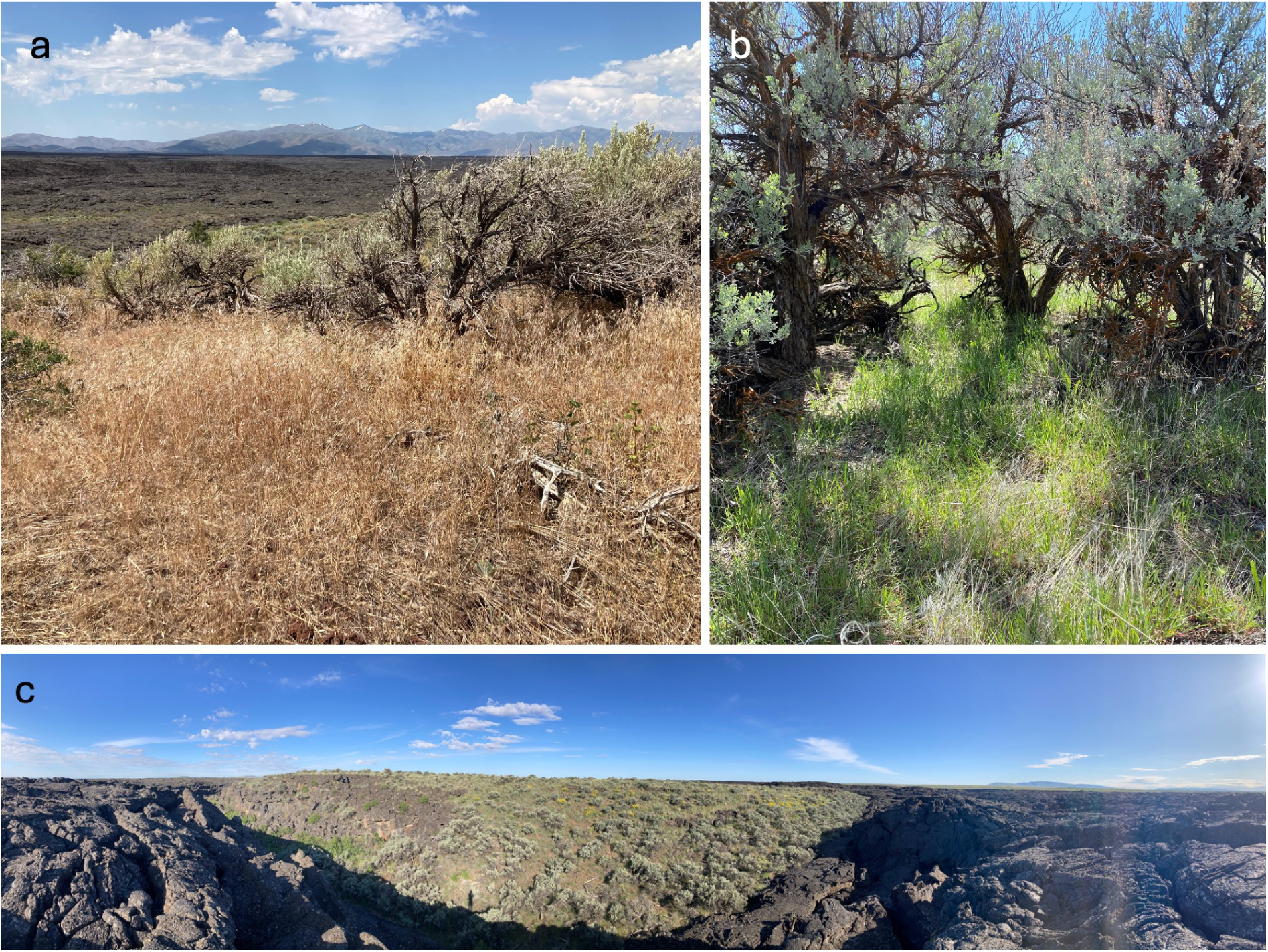
Severe infestations of cheatgrass (*Bromus tectorum* L.) are common among kīpukas in the northern Great Basin. **(a)** a heavily-invaded southwest-facing slope in a kīpuka in the northern portion of Craters of the Moon National Monument and Preserve (CRMO) at approximately 1,675 m (5,500 ft). **(b)** dense cheatgrass understory beneath Wyoming big sagebrush (*Artemisia tridentata* ssp. *wyomingensis*) in a kīpuka isolated by >3 km of lava at approximately 1,370 m (4,500 ft) in CRMO. **(c)** A panoramic view from the edge of a small kīpuka in the Jordan Craters lava flow in eastern Oregon. Photo credits: J. Smith (a & c) and D. Stucki (b).

Cheatgrass (*Bromus tectorum* L.) is almost certainly the species most consistently reflected in our remote sensing data. It was the only invasive annual grass observed in kīpukas during field reconnaissance in 2023 (J. T. Smith, *unpublished data*). Reporting on monitoring efforts in CRMO, Nicolli (2020) called cheatgrass “the most widespread and threatening non-native species in the park,” with several kīpukas showing complete plot-level occupancy. Across seven sampled kīpukas, cheatgrass was recorded in 87% of (1 m^2^) plots and 49% of plots were “heavily infested” with cheatgrass (i.e., >25% cover; Nicolli, 2020). No other invasive annual grasses were reported to be common, and the most prevalent invasive annual forb (*Sisymbrium altissimum* L.) was found in only 11% of plots (Nicolli 2020). Thus, although our remotely sensed data nominally include a larger group of species, we focus primarily on cheatgrass in the remainder of the discussion.

The effects estimated herein represent the average effect of protection from disturbance among the sampled kīpukas. The large number of kīpukas included in the analysis, and their distribution across a broad swath of the northern Great Basin, bolster our confidence that these effects are representative for this region. We caution, however, that extrapolation beyond the sampled population—i.e., to unsampled ecoregions, vegetation communities, or elevations—is not without the usual risks. The kīpukas included in our analysis span a subset of the climatic gradients linked to resistance to invasion in the sagebrush biome, such as temperature, snow-free period, and summer precipitation (Chambers et al. 2014, 2023). Disturbance may have stronger or weaker effects among plant communities occupying different regions of these gradients (Taylor et al. 2014).

Biotic resistance is evidently insufficient to prevent severe invasion and degradation where abiotic conditions are favorable and adequate propagule pressure exists. Biotic interactions (e.g., herbivory, competition, parasitism) commonly limit the severity of infestations but are incapable of completely repelling invaders (Levine et al. 2004), particularly where propagule pressure is high (Von Holle and Simberloff 2005, Colautti et al. 2006). Although propagule pressure was presumably reduced among kīpukas compared to the disturbed sites, cheatgrass has been widespread and abundant in this region for many decades. For example, cheatgrass was recorded in 70% of the AIM field plots used for calibration and validation (235 of 334; see Appendix S1), making it the second most common vascular plant species in this reference dataset. If sufficient to overcome biotic resistance in kīpukas, it is doubtful whether propagule pressure is a limiting factor anywhere in this region. This suggests management strategies focused solely on reducing propagule pressure (e.g., through buffer zones or vector control) may have limited effectiveness once invasion is widespread.

What explains the difference between our findings and studies reporting strong effects of disturbance on cheatgrass demography or abundance? An important distinction lies in the time scales involved. Most experiments and observational studies document short-term responses; removal experiments typically quantify effects of release from competition for 1–2 growing seasons (Beckstead and Augspurger 2004, Chambers et al. 2007), while observational studies infer relationships from spatial co-variation in disturbance and invader abundance captured at a point in time (Reisner et al. 2013, Condon and Pyke 2018, Root et al. 2020). These studies may better reveal how disturbance accelerates invasion in the short run and provide important mechanistic insights, but cannot address longer-term outcomes. In contrast, our approach is insensitive to potentially transient short-term dynamics following disturbance events but provides unique insight into the long-term consequences of altered disturbance regimes. Over time, the distinction between gradual and accelerated invasion becomes increasingly insignificant.

Several traits enable cheatgrass to persist and increase in abundance once established. It builds large and partially persistent soil seed banks even where it appears to be a minor member of plant communities (Hassan and West 1986). Seeds can remain viable for 2–3 years (Hulbert 1955, Young et al. 1969, Smith et al. 2008), allowing populations to survive complete reproductive failure (Rice and Dyer 2001, Baughman et al. 2016). Though emergence is typically concentrated in autumn and secondarily in spring, cheatgrass seeds can germinate at nearly any time of the year (Mack and Pyke 1983, Mack and Pyke 1984) and produce substantial seed rain even at sparse densities, ensuring rapid population recovery when conditions improve (Young and Evans 1978). Germinating at lower temperatures, cheatgrass seedlings emerge earlier than their native competitors (Wilson et al. 1974, Martens et al. 1994) and preemptively deplete limiting soil water, thereby suppressing growth (Melgoza et al. 1990, Nasri and Doescher 1995) and recruitment (Robertson and Pearse 1945, Harris 1967, Rafferty and Young 2002, Humphrey and Schupp 2004) of neighboring perennial bunchgrasses and shrubs.

These traits may interact with natural fluctuations in resources to drive increasing annual grass dominance within undisturbed communities. In drylands like the Great Basin, periodic droughts (Wang et al. 2012) often weaken or kill resident perennials, opening windows of heightened invasibility with the return of normal precipitation (Davis et al. 2000, Chambers et al. 2007). Where it has established seed banks, cheatgrass responds faster than native perennials to improving conditions (Stewart and Hull 1949). Rapid expansions in the area dominated by invasive annual grasses have coincided with drought recoveries in recent decades (Smith et al. 2022), echoing observations following the severe droughts of the 1930s (Pechanec et al. 1937, Stewart and Hull 1949) and 1950s (Tisdale et al. 1965). Plant community dynamics within kīpukas display these same signatures; the abrupt increase in annuals between 2015 and 2018 in CRMO followed a persistent drought that reduced perennial cover (Fig. 5). These dynamics suggest that post-drought recovery periods may represent predictable windows of heightened risk, when targeted early detection and rapid response or strategic seed-bank depletion could most effectively limit cheatgrass spread and impacts to native perennials.

**Figure 5:**
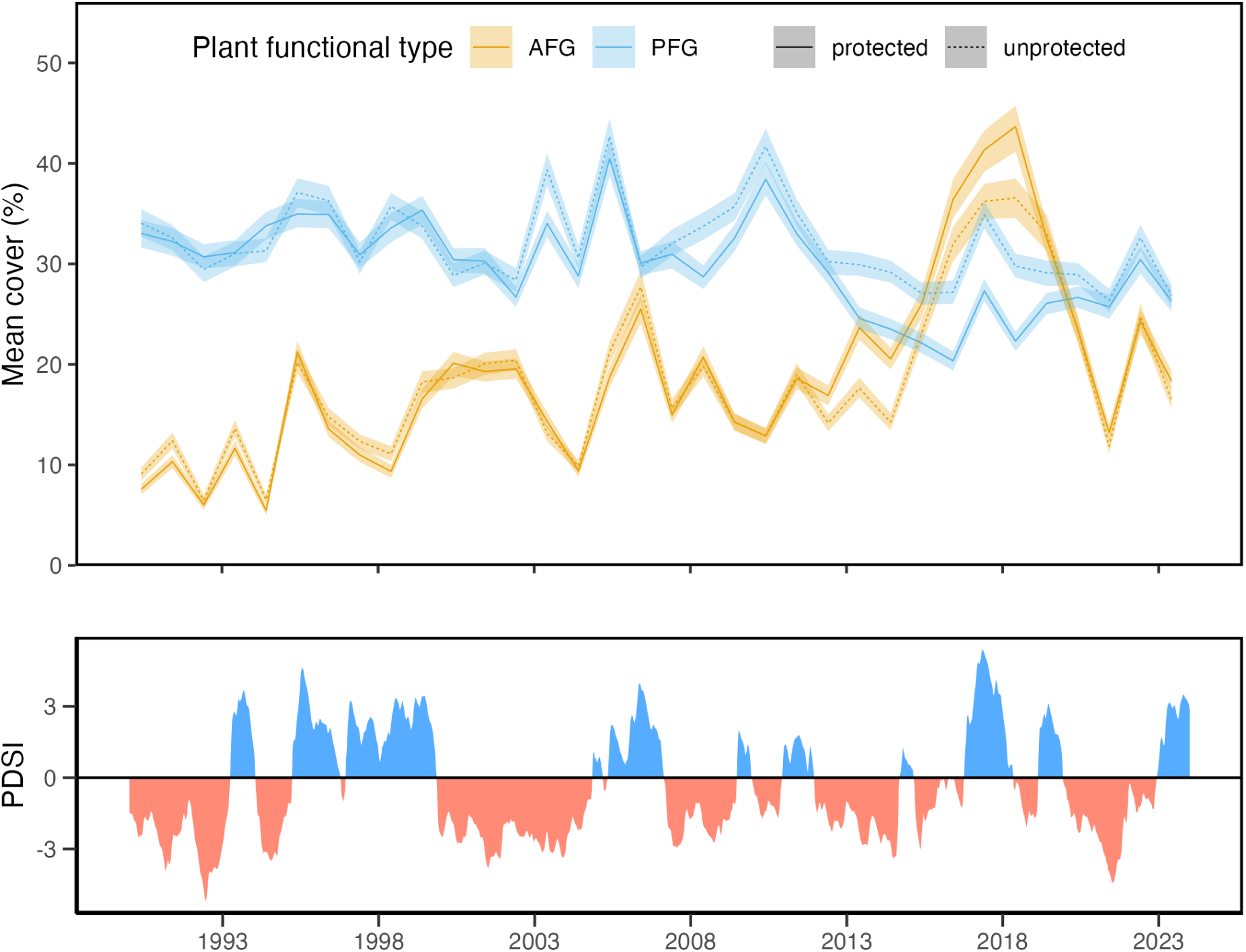
A 34-year time series of cover (mean and 95% confidence interval) of annual forbs and grasses (AFG; in gold) and perennial forbs and grasses (PFG; in blue) among 197 kīpukas in Craters of the Moon National Monument and Preserve in southeastern Idaho and matched unprotected sites (top panel). Data are from the Landsat-based RAP v3 (Allred et al. 2021) rather than the Sentinel-2-based RAP s2 because the latter only extends back to 2018. Protected and unprotected plant communities experienced similar annual fluctuations and long-term trends in these key functional groups, with cover of annuals often reaching new highs with the return of wetter conditions after a drought (average Palmer Drought Severity Index [PDSI] shown in bottom panel).

The rapid transformation of western North American shrublands by invasive annual grasses raises urgent questions about how to conserve what remains (Davies et al. 2021, Doherty et al. 2022). Competing prescriptions reflect different diagnoses of the driver. Some blame excessive fire and advocate doubling down on fire prevention and suppression (Whisenant 1990, Jewell 2015). Others view the worsening invasion as a symptom of more than a century of livestock grazing in an ecosystem too fragile to sustain it and call for the substantial reduction or cessation of grazing (Mack and Thompson 1982, Kauffman et al. 2022). Many point to “all of the above” and suggest passive restoration may be possible with a reduced human footprint on the land (Pyke et al. 2016). Foundational to all of these views is the assumption that invasive annual grasses are passengers of fine-scale disturbance processes cumulatively affecting large areas over time. The unmitigated invasion of kīpukas—the least disturbed sagebrush ecosystems on Earth—demands a critical reevaluation of that assumption.

Decades of debate over the roles of fire and grazing in promoting cheatgrass invasion have overlooked an uncomfortable reality: where abiotic conditions favor invasion, reducing disturbance may slow the pace of ecosystem transformation but is unlikely to meaningfully alter its trajectory. This implies a need to recalibrate conservation priorities in sagebrush ecosystems, where management has for too long been “reactionary and crisis-driven” (Boyd et al. 2024). Halting the cycle of degradation will require a strategic and forward-looking approach that recognizes invasive annual grasses as drivers of change and directly exploits their vulnerabilities (e.g., Sebastian et al. 2017). Without such proactive intervention, large portions of the sagebrush biome will continue their steady conversion into biologically impoverished exotic annual grasslands.

## Acknowledgments

This research was funded by the USDA Agricultural Research Service and the USDA Natural Resources Conservation Service’s Working Lands for Wildlife. Any opinions, findings, conclusions, or recommendations expressed in this publication are those of the authors and do not necessarily reflect the view of the USDA. USDA is an equal opportunity provider and employer. Ray Callaway provided helpful comments that improved the quality of the manuscript, and Jeff Lonneker and Gilbert Moreno provided data and field assistance in Craters of the Moon National Monument and Preserve.

## Notes

https://doi.org/10.5281/zenodo.20332069

### Competing Interest Statement

The authors have declared no competing interest.

## References

1. Abadie, A., and J. Spiess. 2022. Robust post-matching inference. J. Am. Stat. Assoc. 117:983–995.

2. Allred, B. W., B. T. Bestelmeyer, C. S. Boyd, C. Brown, K. W. Davies, M. C. Duniway, L. M. Ellsworth, T. A. Erickson, S. D. Fuhlendorf, T. V. Griffiths, V. Jansen, M. O. Jones, J. Karl, A. Knight, J. D. Maestas, J. J. Maynard, S. E. McCord, D. E. Naugle, H. D. Starns, D. Twidwell, and D. R. Uden. 2021. Improving Landsat predictions of rangeland fractional cover with multitask learning and uncertainty. Methods Ecol. Evol. 12:841–849.

3. Allred, B. W., S. E. McCord, T. J. Assal, B. T. Bestelmeyer, C. S. Boyd, A. C. Brooks, S. M. Cady, M. C. Duniway, S. D. Fuhlendorf, S. A. Green, G. R. Harrison, E. R. Jensen, E. J. Kachergis, A. C. Knight, C. M. Mattilio, B. A. Mealor, D. E. Naugle, D. O’Leary, P. J. Olsoy, E. S. Peirce, J. R. Reinhardt, R. K. Shriver, J. T. Smith, J. D. Tack, A. M. Tanner, E. P. Tanner, D. Twidwell, N. P. Webb, and S. L. Morford. 2025. Sentinel-2 based estimates of rangeland fractional cover and canopy gap class for the western United States. Sci. Data 12:1889.

4. Arel-Bundock, V., N. Greifer, and A. Heiss. 2024. How to interpret statistical models using marginaleffects for R and python. J. Stat. Softw. 111:1–32.

5. Bangert, R., and N. Huntly. 2010. The distribution of native and exotic plants in a naturally fragmented sagebrush-steppe landscape. Biol. Invasions 12:1627–1640.

6. Bansal, S., and R. L. Sheley. 2016. Annual grass invasion in sagebrush steppe: The relative importance of climate, soil properties and biotic interactions. Oecologia 181:543–557.

7. Bates, D., M. Maechler, B. Bolker, and S. Walker. 2015. Fitting linear mixed-effects models using lme4. J. Stat. Softw. 67:1–48.

8. Bauer, J. T. 2012. Invasive species: “Back-seat drivers” of ecosystem change? Biol. Invasions 14:1295–1304.

9. Baughman, O. W., S. E. Meyer, Z. T. Aanderud, and E. A. Leger. 2016. Cheatgrass die-offs as an opportunity for restoration in the Great Basin, USA: Will local or commercial native plants succeed where exotic invaders fail? J. Arid Environ. 124:193–204.

10. Beckstead, J., and C. K. Augspurger. 2004. An experimental test of resistance to cheatgrass invasion: Limiting resources at different life stages. Biol. Invasions 6:417–432.

11. Boyd, C. S., M. K. Creutzburg, A. V. Kumar, J. T. Smith, K. E. Doherty, B. A. Mealor, J. B. Bradford, M. Cahill, S. M. Copeland, C. A. Duquette, L. Garner, M. C. Holdrege, B. Sparklin, and T. B. Cross. 2024. A strategic and science-based framework for management of invasive annual grasses in the sagebrush biome. Rangeland Ecol. Manage. 97:61–72.

12. Brooks, M. L., D. A. Pyke, K. E. M. Galley, and T. P. Wilson. 2001. Invasive plants and fire in the deserts of North America. Proceedings of the invasive species workshop: The role of fire in the control and spread of invasive species, eds galley KEM, wilson TP (tall timbers research station, tallahassee, FL). firescience.gov.

13. Brown, C. F., M. R. Kazmierski, V. J. Pasquarella, W. J. Rucklidge, M. Samsikova, C. Zhang, E. Shelhamer, E. Lahera, O. Wiles, S. Ilyushchenko, N. Gorelick, L. L. Zhang, S. Alj, E. Schechter, S. Askay, O. Guinan, R. Moore, A. Boukouvalas, and P. Kohli. 2025. AlphaEarth foundations: An embedding field model for accurate and efficient global mapping from sparse label data. arXiv [cs.CV].

14. Callaway, R. M., and W. M. Ridenour. 2004. Novel weapons: Invasive success and the evolution of increased competitive ability. Front. Ecol. Environ. 2:436–443.

15. Carneiro, L., N. O. R. Miiller, R. N. Cuthbert, and J. R. S. Vitule. 2024. Biological invasions negatively impact global protected areas. Sci. Total Environ. 948:174823.

16. Catford, J. A., C. C. Daehler, H. T. Murphy, A. W. Sheppard, B. D. Hardesty, D. A. Westcott, M. Rejmánek, P. J. Bellingham, J. Pergl, C. C. Horvitz, and P. E. Hulme. 2012. The intermediate disturbance hypothesis and plant invasions: Implications for species richness and management. Perspect. Plant Ecol. Evol. Syst. 14:231–241.

17. Chambers, J. C., B. A. Bradley, C. S. Brown, C. D’Antonio, M. J. Germino, J. B. Grace, S. P. Hardegree, R. F. Miller, and D. A. Pyke. 2014. Resilience to stress and disturbance, and resistance to *Bromus tectorum* L. Invasion in cold desert shrublands of western north America. Ecosystems 17:360–375.

18. Chambers, J. C., J. L. Brown, J. B. Bradford, David I. Board, S. B. Campbell, K. J. Clause, B. Hanberry, D. R. Schlaepfer, and A. K. Urza. 2023. New indicators of ecological resilience and invasion resistance to support prioritization and management in the sagebrush biome, United States. Front. Ecol. Evol. 10:1009268.

19. Chambers, J. C., B. A. Roundy, R. R. Blank, S. E. Meyer, and A. Whittaker. 2007. What makes Great Basin sagebrush ecosystems invasible by *Bromus tectorum*? Ecol. Monogr. 77:117–145.

20. Coates, P. S., M. A. Ricca, B. G. Prochazka, M. L. Brooks, K. E. Doherty, T. Kroger, E. J. Blomberg, C. A. Hagen, and M. L. Casazza. 2016. Wildfire, climate, and invasive grass interactions negatively impact an indicator species by reshaping sagebrush ecosystems. Proceedings of the National Academy of Sciences 113:12745–12750.

21. Colautti, R. I., I. A. Grigorovich, and H. J. MacIsaac. 2006. Propagule pressure: A null model for biological invasions. Biol. Invasions 8:1023–1037.

22. Condon, L. A., and D. A. Pyke. 2018. Fire and grazing influence site resistance to *Bromus tectorum* through their effects on shrub, bunchgrass and biocrust communities in the Great Basin (USA). Ecosystems 21:1416–1431.

23. Copeland, S. M., D. L. Hoover, D. J. Augustine, J. D. Bates, C. S. Boyd, K. W. Davies, J. D. Derner, M. C. Duniway, L. M. Porensky, and L. T. Vermeire. 2023. Variable effects of long-term livestock grazing across the western United States suggest diverse approaches are needed to meet global change challenges. Appl. Veg. Sci. 26:e12719.

24. D’Antonio, C. M., and P. M. Vitousek. 1992. Biological invasions by exotic grasses, the grass/fire cycle, and global change. Annu. Rev. Ecol. Syst. 23:63–87.

25. Daubenmire, R. F. 1940. Plant succession due to overgrazing in the Agropyron bunchgrass prairie of southeastern Washington. Ecology 21:55–64.

26. Davies, K. W., E. A. Leger, C. S. Boyd, and L. M. Hallett. 2021. Living with exotic annual grasses in the sagebrush ecosystem. J. Environ. Manage. 288:112417.

27. Davies, K. W., T. J. Svejcar, and J. D. Bates. 2009. Interaction of historical and nonhistorical disturbances maintains native plant communities. Ecol. Appl. 19:1536–1545.

28. Davis, M. A., J. P. Grime, and K. Thompson. 2000. Fluctuating resources in plant communities: A general theory of invasibility. J. Ecol. 88:528–534.

29. Doherty, K., D. M. Theobald, J. B. Bradford, L. A. Wiechman, G. Bedrosian, C. S. Boyd, M. Cahill, P. S. Coates, M. K. Creutzburg, M. R. Crist, S. P. Finn, A. V. Kumar, C. E. Littlefield, J. D. Maestas, K. L. Prentice, B. G. Prochazka, T. E. Remington, W. D. Sparklin, J. C. Tull, Z. Wurtzebach, and K. A. Zeller. 2022. A sagebrush conservation design to proactively restore America’s sagebrush biome. U.S. Geological Survey.

30. Driscoll, R. S. 1964. A relict area in the central Oregon juniper zone. Ecology 45:345–353.

31. Eidenshink, J., B. Schwind, K. Brewer, Z.-L. Zhu, B. Quayle, and S. Howard. 2007. A project for monitoring trends in burn severity. Fire Ecology 3:3–21.

32. Elton, C. S. 1958. The ecology of invasions by animals and plants. Methuen, London.

33. Foxcroft, L. C., P. Pyšek, D. M. Richardson, P. Genovesi, and S. MacFadyen. 2017. Plant invasion science in protected areas: Progress and priorities. Biol. Invasions 19:1353–1378.

34. Fusco, E. J., J. T. Finn, J. K. Balch, R. C. Nagy, and B. A. Bradley. 2019. Invasive grasses increase fire occurrence and frequency across US ecoregions. Proc. Natl. Acad. Sci. U. S. A. 116:23594–23599.

35. Gale, S., and N. Goodrich. 1999. Cheatgrass frequency at two relic sites within the pinyon-juniper belt of red canyon. *in* S. B. Monsen and R. Stevens, editors. Proceedings: Ecology and management of pinyon-juniper communities within the interior west. U.S. Department of Agriculture, Forest Service, Rocky Mountain Research Station.

36. Germino, M. J., J. Belnap, J. M. Stark, E. B. Allen, and B. M. Rau. 2016. Ecosystem impacts of exotic annual invaders in the genus Bromus. Pages 61–95 in M. J. Germino, J. C. Chambers, and C. S. Brown, editors. Exotic brome-grasses in arid and semiarid ecosystems of the western US: Causes, consequences, and management implications. Springer International Publishing, Cham.

37. Harris, G. A. 1967. Some competitive relationships between *Agropyron spicatum* and *Bromus tectorum*. Ecol. Monogr. 37:89–111.

38. Hassan, M. A., and N. E. West. 1986. Dynamics of soil seeds pools in burned and unburned sagebrush semi-deserts. Ecology 67:269–272.

39. Hierro, J. L., D. Villarreal, Ö. Eren, J. M. Graham, and R. M. Callaway. 2006. Disturbance facilitates invasion: The effects are stronger abroad than at home. The American Naturalist 168:144–156.

40. Hille Ris Lambers, J., S. G. Yelenik, B. P. Colman, and J. M. Levine. 2010. California annual grass invaders: The drivers or passengers of change?: Annual grass invaders in California. J. Ecol. 98:1147–1156.

41. Ho, D. E., K. Imai, G. King, and E. A. Stuart. 2007. Matching as nonparametric preprocessing for reducing model dependence in parametric causal inference. Polit. Anal. 15:199–236.

42. Ho, D. E., K. Imai, G. King, and E. A. Stuart. 2011. MatchIt: Nonparametric preprocessing for parametric causal inference. J. Stat. Softw. 42:1–28.

43. Hobart, B. K., W. E. Moss, M. C. Cook, R. C. Nagy, and V. J. McKenzie. 2026. Annual grass invasion is transforming the sagebrush biome’s songbird communities. Front. Ecol. Environ.

44. Hobbs, R. J., and L. F. Huenneke. 1992. Disturbance, diversity, and invasion: Implications for conservation. Conserv. Biol. 6:324–337.

45. Hooker, T. D., J. M. Stark, U. Norton, A. Joshua Leffler, M. Peek, and R. Ryel. 2008. Distribution of ecosystem C and N within contrasting vegetation types in a semiarid rangeland in the Great Basin, USA. Biogeochemistry 90:291–308.

46. Hulbert, L. C. 1955. Ecological studies of *Bromus tectorum* and other annual bromegrasses. Ecol. Monogr. 25:181–213.

47. Humphrey, L. D., and E. W. Schupp. 2004. Competition as a barrier to establishment of a native perennial grass (*Elymus elymoides*) in alien annual grass (*Bromus tectorum*) communities. J. Arid Environ. 58:405–422.

48. Jauni, M., S. Gripenberg, and S. Ramula. 2015. Non-native plant species benefit from disturbance: A meta-analysis. Oikos 124:122–129.

49. Jewell, S. 2015. Order no. 3336: Rangeland fire prevention, management and restoration. Department of Interior.

50. Kauffman, J. B., R. L. Beschta, P. M. Lacy, and M. Liverman. 2022. Livestock use on public lands in the western USA exacerbates climate change: Implications for climate change mitigation and adaptation. Environ. Manage. 69:1137–1152.

51. Kindschy, R. R. 1994. Pristine vegetation of the jordan crater kipukas: 1978-91. *in* S. B. Monsen and S. G. Ketchum, editors. General technical report INT-GTR-313: Proceedings—ecology and management of annual rangelands. U.S. Department of Agriculture, Forest Service, Intermountain Research Station.

52. Leffler, A. J., T. A. Monaco, J. J. James, and R. L. Sheley. 2016. Importance of soil and plant community disturbance for establishment of *Bromus tectorum* in the intermountain west, USA. NeoBiota 30:111–125.

53. Leopold, A. 1941. Cheat takes over. The Land.

54. Levine, J. M., P. B. Adler, and S. G. Yelenik. 2004. A meta-analysis of biotic resistance to exotic plant invasions: Biotic resistance to plant invasion. Ecol. Lett. 7:975–989.

55. MacDougall, A. S., and R. Turkington. 2005. Are invasive species the drivers or passengers of change in degraded ecosystems? Ecology 86:42–55.

56. Mack, R. N. 1981. Invasion of *Bromus tectorum* L. Into western north America: An ecological chronicle. Agro-Ecosyst. 7:145–165.

57. Mack, R. N., and D. A. Pyke. 1983. The demography of *Bromus tectorum*: Variation in time and space. J. Ecol. 71:69.

58. Mack, R. N., and D. A. Pyke. 1984. The demography of *Bromus tectorum*: The role of microclimate, grazing and disease. J. Ecol. 72:731–748.

59. Mack, R. N., and J. N. Thompson. 1982. Evolution in steppe with few large, hooved mammals. Am. Nat. 119:757–773.

60. Mahood, A. L., and J. K. Balch. 2019. Repeated fires reduce plant diversity in low-elevation Wyoming big sagebrush ecosystems (1984–2014). Ecosphere 10:e02591.

61. Martens, E., D. Palmquist, J. A. Young, S. B. Monsen, and S. G. Kitchen. 1994. Temperature profiles for germination of cheatgrass versus native perennial bunchgrasses. Proceedings—Ecology and management of annual rangelands (SB Monsen and SG Kitchen, editors). US Department of Agriculture Forest Service, Intermountain Research Station, General Technical Report INT-GTR-313:238–243.

62. Melgoza, G., R. S. Nowak, and R. J. Tausch. 1990. Soil water exploitation after fire: Competition between *Bromus tectorum* (cheatgrass) and two native species. Oecologia 83:7–13.

63. Nasri, M., and P. S. Doescher. 1995. Effect of competition by cheatgrass on shoot growth of Idaho fescue. Rangeland Ecol. Manage. 48:402–405.

64. Nauman, T. W., S. Kienast-Brown, S. M. Roecker, C. Brungard, D. White, J. Philippe, and J. A. Thompson. 2024. Soil landscapes of the United States (SOLUS): Developing predictive soil property maps of the conterminous United States using hybrid training sets. Soil Sci. Soc. Am. J. 88:2046–2065.

65. Nicolli, M. M. 2020. Sagebrush steppe vegetation monitoring in Craters of the Moon national monument and preserve: 2019 annual report. National Park Service.

66. Omernik, J. M., and G. E. Griffith. 2014. Ecoregions of the conterminous United States: Evolution of a hierarchical spatial framework. Environ. Manage. 54:1249–1266.

67. Passey, H. B., and V. K. Hugie. 1963. Some plant-soil relationships on an ungrazed range area of southeastern Idaho. J. Range Manage. 16:113.

68. Passey, H. B., V. K. Hugie, E. W. Williams, and D. E. Ball. 1982. Relationships between soil, plant community, and climate on rangelands of the intermountain west. US Department of Agriculture, Soil Conservation Service.

69. Pearson, D. E., Y. K. Ortega, Ö. Eren, D. Villarreal, Y. Lekberg, and J. Hierro. 2023. Exotic success following disturbance explained by weak native resilience and ruderal exotic bias. J. Ecol. 111:2412–2423.

70. Pechanec, J. F., G. D. Pickford, and G. Stewart. 1937. Effects of the 1934 drought on native vegetation of the upper snake river plans, Idaho. Ecology 18:490–505.

71. Porensky, L. M., R. McGee, and D. W. Pellatz. 2020. Long-term grazing removal increased invasion and reduced native plant abundance and diversity in a sagebrush grassland. Glob. Ecol. Conserv. 24:e01267.

72. Proctor, J., T. Carleton, and S. Sum. 2023. Parameter recovery using remotely sensed variables. National Bureau of Economic Research.

73. Pyke, D. A., J. C. Chambers, J. L. Beck, M. L. Brooks, and B. A. Mealor. 2016. Land uses, fire, and invasion: Exotic annual Bromus and human dimensions. Pages 307–337 in M. J. Germino, J. C. Chambers, and C. S. Brown, editors. Exotic brome-grasses in arid and semiarid ecosystems of the western US: Causes, consequences, and management implications. Springer International Publishing, Cham.

74. Rafferty, D. L., and J. A. Young. 2002. Cheatgrass competition and establishment of desert needlegrass seedlings. Rangeland Ecol. Manage. 55:70–72.

75. Reisner, M. D., J. B. Grace, D. A. Pyke, and P. S. Doescher. 2013. Conditions favouring *Bromus tectorum* dominance of endangered sagebrush steppe ecosystems. J. Appl. Ecol. 50:1039–1049.

76. Ren, J., J. Chen, C. Xu, J. van de Koppel, M. S. Thomsen, S. Qiu, F. Cheng, W. Song, Q.-X. Liu, C. Xu, J. Bai, Y. Zhang, B. Cui, M. D. Bertness, B. R. Silliman, B. Li, and Q. He. 2021. An invasive species erodes the performance of coastal wetland protected areas. Sci. Adv. 7:eabi8943.

77. Rice, K. J., and A. R. Dyer. 2001. Seed aging, delayed germination and reduced competitive ability in *Bromus tectorum*. Plant Ecol. 155:237–243.

78. Rinella, M. J., L. T. Vermeire, and J. P. Angerer. 2024. Integrating experiments and monitoring reveals extreme sensitivity of invasive winter annuals to precipitation. Ecol. Appl. 34:e3051.

79. Robertson, J. H., and C. K. Pearse. 1945. Artificial reseeding and the closed community. Northwest Sci. 19:58–66.

80. Root, H. T., J. E. D. Miller, and R. Rosentreter. 2020. Grazing disturbance promotes exotic annual grasses by degrading soil biocrust communities. Ecol. Appl. 30:e02016.

81. Rubin, D. B. 1974. Estimating causal effects of treatments in randomized and nonrandomized studies. J. Educ. Psychol. 66:688–701.

82. Rubin, D. B. 2001. Using propensity scores to help design observational studies: Application to the tobacco litigation. Health Serv. Outcomes Res. Methodol. 2:169–188.

83. Schaeffer, S. M., S. E. Ziegler, J. Belnap, and R. D. Evans. 2012. Effects of *Bromus tectorum* invasion on microbial carbon and nitrogen cycling in two adjacent undisturbed arid grassland communities. Biogeochemistry 111:427–441.

84. Sebastian, D. J., S. J. Nissen, J. R. Sebastian, and K. G. Beck. 2017. Seed bank depletion: The key to long-term downy brome (*Bromus tectorum* L.) management. Rangeland Ecol. Manage. 70:477–483.

85. Shackleton, R. T., L. C. Foxcroft, P. Pyšek, L. E. Wood, and D. M. Richardson. 2020. Assessing biological invasions in protected areas after 30 years: Revisiting nature reserves targeted by the 1980s SCOPE programme. Biol. Conserv. 243:108424.

86. Smith, D. C., S. E. Meyer, and V. J. Anderson. 2008. Factors affecting *Bromus tectorum* seed bank carryover in western Utah. Rangeland Ecol. Manage. 61:430–436.

87. Smith, J. T., B. W. Allred, C. S. Boyd, K. W. Davies, M. O. Jones, A. R. Kleinhesselink, J. D. Maestas, S. L. Morford, and D. E. Naugle. 2022. The elevational ascent and spread of exotic annual grass dominance in the Great Basin, USA. Divers. Distrib. 28:83–96.

88. Smith, J. T., B. W. Allred, C. S. Boyd, K. W. Davies, A. R. Kleinhesselink, S. L. Morford, and D. E. Naugle. 2023. Fire needs annual grasses more than annual grasses need fire. Biol. Conserv. 286:110299.

89. Sperry, L. J., J. Belnap, and R. D. Evans. 2006. *Bromus tectorum* invasion alters nitrogen dynamics in an undisturbed arid grassland ecosystem. Ecology 87:603–615.

90. Splawa-Neyman, J., D. M. Dabrowska, and T. P. Speed. 1990. On the application of probability theory to agricultural experiments. Essay on principles. Section 9. Stat. Sci. 5:465–472.

91. Stewart, G., and A. C. Hull. 1949. Cheatgrass (*Bromus tectorum* L.)—an ecologic intruder in southern Idaho. Ecology 30:58–74.

92. Stuart, E. A. 2010. Matching methods for causal inference: A review and a look forward. Stat. Sci. 25:1–21.

93. Tausch, R. J., T. Svejcar, and J. W. Burkhardt. 1994. Patterns of annual grass dominance on anaho island: Implications for Great Basin vegetation management. Pages 120–125 *in* S. B. Monsen and S. G. Ketchum, editors. General technical report INT-GTR-313: Proceedings—ecology and management of annual rangelands. U.S. Department of Agriculture, Forest Service, Intermountain Research Station.

94. Taylor, K., T. Brummer, L. J. Rew, M. Lavin, and B. D. Maxwell. 2014. *Bromus tectorum* response to fire varies with climate conditions. Ecosystems 17:960–973.

95. Theobald, D. M., D. Harrison-Atlas, W. B. Monahan, and C. M. Albano. 2015. Ecologically-relevant maps of landforms and physiographic diversity for climate adaptation planning. PLoS One 10:e0143619.

96. Tilman, D. 1994. Competition and biodiversity in spatially structured habitats. Ecology 75:2–16.

97. Tisdale, E. W., M. Hironaka, and M. A. Fosberg. 1965. An area of pristine vegetation in craters of the moon national monument, Idaho. Ecology 46:349–352.

98. Urza, A. K., D. I. Board, J. B. Bradford, J. L. Brown, J. C. Chambers, D. R. Schlaepfer, and K. C. Short. 2024. Disentangling drivers of annual grass invasion: Abiotic susceptibility vs. Fire-induced conversion to cheatgrass dominance in the sagebrush biome. Biol. Conserv. 297:110737.

99. Vitousek, P. M., L. R. Walker, L. D. Whiteaker, D. Mueller-Dombois, and P. A. Matson. 1987. Biological invasion by Myrica faya alters ecosystem development in Hawaii. Science 238:802–804.

100. Von Holle, B., and D. Simberloff. 2005. Ecological resistance to biological invasion overwhelmed by propagule pressure. Ecology 86:3212–3218.

101. Wang, S.-Y., R. R. Gillies, and T. Reichler. 2012. Multidecadal drought cycles in the Great Basin recorded by the great salt lake: Modulation from a transition-phase teleconnection. J. Clim. 25:1711–1721.

102. Wauchope, H. S., J. P. G. Jones, J. Geldmann, B. I. Simmons, T. Amano, D. E. Blanco, R. A. Fuller, A. Johnston, T. Langendoen, T. Mundkur, S. Nagy, and W. J. Sutherland. 2022. Protected areas have a mixed impact on waterbirds, but management helps. Nature 605:103–107.

103. Whisenant, S. G. 1990. Changing fire frequencies on Idaho’s snake river plains: Ecological and management implications. USDA Forest Service, Ogden, UT.

104. Williamson, M. A., E. Fleishman, R. C. Mac Nally, J. C. Chambers, B. A. Bradley, D. S. Dobkin, David I. Board, F. A. Fogarty, N. Horning, M. Leu, and M. Wohlfeil Zillig. 2020. Fire, livestock grazing, topography, and precipitation affect occurrence and prevalence of cheatgrass (*Bromus tectorum*) in the central Great Basin, USA. Biol. Invasions 22:663–680.

105. Wilson, A. M., D. E. Wondercheck, and C. J. Goebel. 1974. Responses of range grass seeds to winter environments. Rangeland Ecol. Manage. 27:120–122.

106. Young, J. A., and R. A. Evans. 1978. Population dynamics after wildfires in sagebrush grasslands. Rangeland Ecol. Manage. 31:283–289.

107. Young, J. A., R. A. Evans, and R. E. Eckert Jr. 1969. Population dynamics of downy brome. Weed Sci. 17:20–26.

108. Young, J. A., R. A. Evans, and J. Major. 1972. Alien plants in the Great Basin. Rangeland Ecol. Manage. 25:194–201.

