## Appendix S1 for "Exclusion of fire and grazing does not reduce annual grass invasion in a sagebrush steppe ecosystem"

**Appendix S1: Regional RAP validation**

Supporting Information for: Exclusion of fire and grazing does not reduce annual grass invasion in a sagebrush steppe ecosystem

Joseph T. Smith, Brady W. Allred, Chad S. Boyd, Kirk W. Davies, Scott L. Morford, David E. Naugle, Thomas J. Rodhouse, and Devin S. Stucki

#### Introduction and methods

Remotely sensed fractional cover datasets such as the Rangeland Analysis Platform (RAP; Allred et al. 2025) mark a generational breakthrough in rangeland monitoring (Jones et al. 2020), overcoming many of the limitations of spatially and temporally sparse and inconsistent field monitoring. However, this progress comes with the tradeoff of reduced taxonomic resolution. These datasets estimate cover of broad functional groups, leaving the species responsible for observed patterns ambiguous. For example, previous studies have used RAP's annual forb and grass (AFG) cover as a proxy for invasive annual grasses in sagebrush ecosystems (Smith et al. 2022, 2023, Boyd et al. 2024). AFG, however, nominally includes many species that are neither invasive nor grasses. Because of the importance of these species in the conservation and management of western North American rangelands, the newer Sentinel-2-based RAP 10 m cover product includes a prediction of invasive annual grass (IAG) cover in addition to AFG. Here, we sought to characterize the regional agreement between the remotely sensed predictions of IAG, perennial forbs and grasses (PFG), shrubs (SHR) and bare ground (BGR) and ground-based estimates of the same variables, as well as verify that IAG is sensitive only to the prevalent invasive annual grass species within our study region.

We extracted RAP cover estimates and summarized species-level line point intercept data at 334 Bureau of Land Management Assessment, Inventory, and Monitoring (BLM-AIM Taylor et al. 2014) plots located within a polygon bounding our matched samples and visited between 2018 and 2023. These plots were selected because they were not used to train the RAP model (Allred et al. 2025). For each functional group, we calculated mean absolute error (MAE) between the RAP-predicted and field-estimated cover. We also calculated the coefficient of determination between RAP-predicted IAG cover and measured cover of the 50 most common species to identify those most strongly associated with high IAG values.

#### Results

Mean absolute error among these 334 BLM-AIM plots was 7.2%, 11.4%, 6.8%, and 6.6% for IAG, PFG, SHR, and BGR, respectively. Coefficients of determination were 0.60, 0.31, 0.47, and 0.70 for IAG, PFG, SHR, and BGR, respectively (Fig. 1).

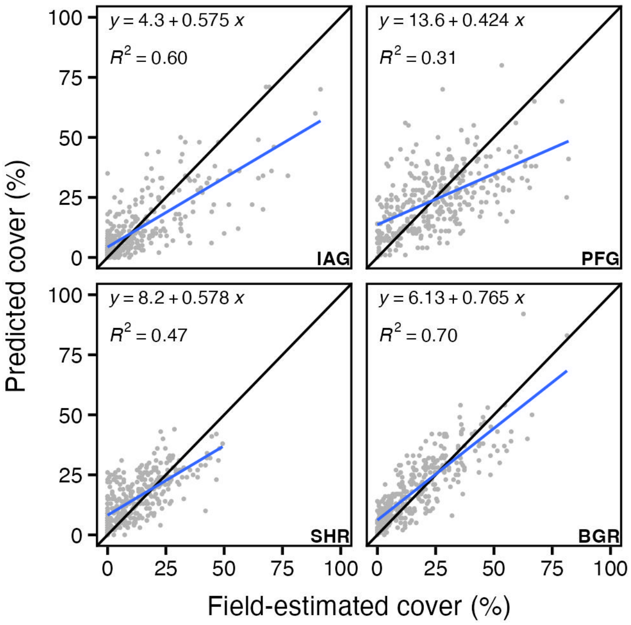


**Figure 1:** Scatterplots showing field-estimated cover (x-axis) vs remotely-sensed cover from the Rangeland Analysis Platform (RAP; y-axis) at 334 independent Bureau of Land Management Assessment, Inventory, and Monitoring (AIM) plots measured within the study area between 2018 and 2023.

Seven invasive annual grass species were recorded among the 334 AIM plots. Cover of the most common invasive annual grass species, *Bromus tectorum* L. (recorded in 70% of plots), explained 39% of the variation in IAG and the second most common invasive annual grass species, *Taeniatherum caput-medusae* (recorded in 12% of plots), explained 25% of the variation in IAG (Fig. S1.2). The remaining 5 species were less regionally prevalent; for example, the third most prevalent invasive annual grass species, *Bromus arvensis* (BRAR5), was only found in 19 plots and generally occurred at low abundance (median cover at occupied plots = 2.97%). The combined cover of all invasive annual grasses explained 60% of the variation in IAG, and no non-target species explained >5% of the variation in IAG (Fig. 2).

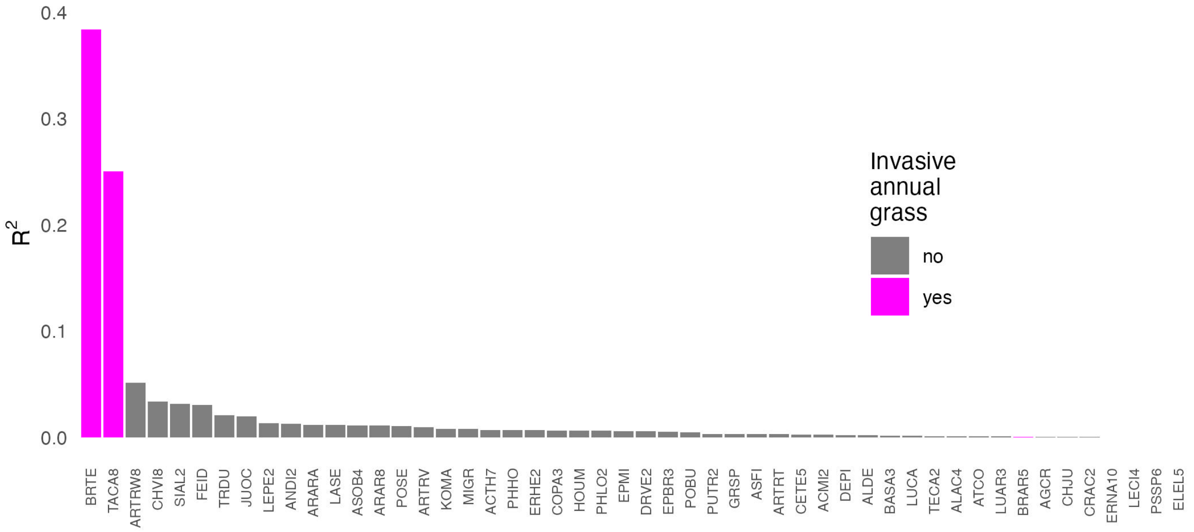


**Figure 2:** Bivariate coefficients of determination ($R^2$) between remotely-sensed estimates of invasive annual grass (IAG) cover from the Rangeland Analysis Platform and field-estimated cover of the 50 most prevalent vascular plant species among 334 independent BLM-AIM monitoring plots measured within the study area since 2018. Only *Bromus tectorum* (BRTE) and *Taeniatherum caput-medusae* (TACA8), both regionally common invasive annual grasses, explained significant variation in IAG.

### References


Allred, B. W., S. E. McCord, T. J. Assal, B. T. Bestelmeyer, C. S. Boyd, A. C. Brooks, S. M. Cady, M. C. Duniway, S. D. Fuhlendorf, S. A. Green, G. R. Harrison, E. R. Jensen, E. J. Kachergis, A. C. Knight, C. M. Mattilio, B. A. Mealor, D. E. Naugle, D. O'Leary, P. J. Olsoy, E. S. Peirce, J. R. Reinhardt, R. K. Shriver, J. T. Smith, J. D. Tack, A. M. Tanner, E. P. Tanner, D. Twidwell, N. P. Webb, and S. L. Morford. 2025. Sentinel-2 based estimates of rangeland fractional cover and canopy gap class for the western United States. Sci. Data 12:1889.

Boyd, C. S., M. K. Creutzburg, A. V. Kumar, J. T. Smith, K. E. Doherty, B. A. Mealor, J. B. Bradford, M. Cahill, S. M. Copeland, C. A. Duquette, L. Garner, M. C. Holdrege, B. Sparklin, and T. B. Cross. 2024. A strategic and science-based framework for management of invasive annual grasses in the sagebrush biome. Rangeland Ecol. Manage. 97:61–72.

Jones, M. O., D. E. Naugle, D. Twidwell, D. R. Uden, J. D. Maestas, and B. W. Allred. 2020. Beyond inventories: Emergence of a new era in rangeland monitoring. Rangeland Ecol. Manage. 73:577–583.

Smith, J. T., B. W. Allred, C. S. Boyd, K. W. Davies, M. O. Jones, A. R. Kleinhesselink, J. D. Maestas, S. L. Morford, and D. E. Naugle. 2022. The elevational ascent and spread of exotic annual grass dominance in the Great Basin, USA. Divers. Distrib. 28:83–96.

Smith, J. T., B. W. Allred, C. S. Boyd, K. W. Davies, A. R. Kleinhesselink, S. L. Morford, and D. E. Naugle. 2023. Fire needs annual grasses more than annual grasses need fire. Biol. Conserv. 286:110299.

Taylor, J. M., E. Kachergis, G. R. Toevs, J. W. Karl, M. R. Bobo, M. G. Karl, S. W. Miller, and C. Spurrier. 2014. AIM-monitoring: A component of the BLM assessment, inventory, and monitoring strategy. U.S. Department of Interior, Bureau of Land Mangement, National Operations Center; ars.usda.gov, Denver, CO.
