## Appendix S2 for "Exclusion of fire and grazing does not reduce annual grass invasion in a sagebrush steppe ecosystem"

**Appendix S2: Evaluation of alternative fractional cover data sources**

Supporting Information for: Exclusion of fire and grazing does not reduce annual grass invasion in a sagebrush steppe ecosystem

Joseph T. Smith, Brady W. Allred, Chad S. Boyd, Kirk W. Davies, Scott L. Morford, David E. Naugle, Thomas J. Rodhouse, and Devin S. Stucki

#### Introduction and methods

Fractional vegetation cover estimation is a new and rapidly evolving branch of remote sensing. Three operational datasets currently provide annual estimates of rangeland vegetation cover in the US: the Rangeland Analysis Platform (RAP; Allred et al. 2021, 2025, Jones et al. 2021), the Rangeland Condition Monitoring Assessment and Projection dataset (RCMAP v7; Rigge et al. 2020, 2021), and the Exotic Annual Grass cover dataset (USGS-EAG; Dahal et al. 2022). Because these datasets were developed independently using different machine learning algorithms and inputs, they commonly predict different values of cover at the same site, reflecting the uncertainties inherent in remote sensing of vegetation.

Our analysis used predictions from the Sentinel-2-based RAP 10 m cover product (Allred et al. 2025), focusing primarily on cover of invasive annual grasses (IAG). To explore the sensitivity of our results to the choice of dataset, we also extracted cover predictions from each of the other two datasets, RCMAP and USGS-EAG, at our matched samples and repeated our analysis. We compared IAG from RAP to rangeland annual herbaceous cover (hereafter RAH) from RCMAP and total exotic annual grass cover (hereafter EAG) from USGS-EAG.

#### Results

Focusing on Set 1, the effect of protection from disturbance on rangeland annual herbaceous (RAH) cover from RCMAP (0.87%, 95% CI from -2.61 to 2.05%) and exotic annual grass cover (EAG) from the USGS product (0.63%, 95% CI from -0.47 to 1.73%) cover were smaller (i.e., less positive) than the effect on invasive annual grass (IAG) cover from RAP (1.51%, 95% CI from 0.37 to 2.64%; Fig. 1). Effect sizes were also smaller for Set 2, with the sign of the effect switching to negative for RAH from RCMAP v7. All confidence intervals overlapped zero for the alternative datasets. In terms of their agreement with the regional field calibration dataset (n = 334; see Appendix S1), IAG had the highest agreement with field estimates of invasive annual grass cover (MAE = 7.2%, R2 = 0.60), followed by RAH (MAE = 9.9%, R2 = 0.42) and EAG (MAE = 10.8%, R2 = 0.30).

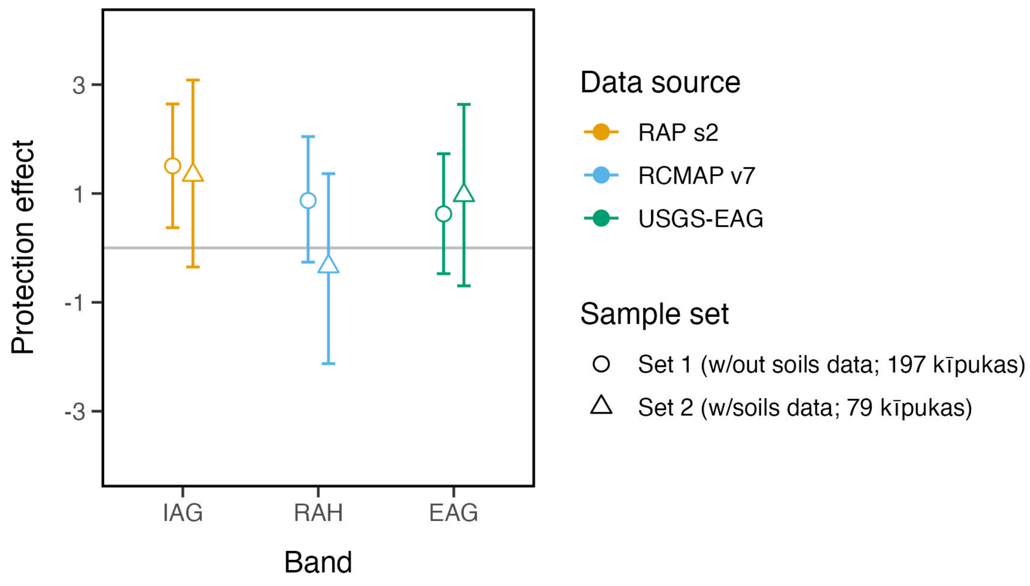


**Figure 1:** Estimated average effect of protection from disturbance (and 95% confidence intervals) on invasive annual grass cover in sagebrush plant communities of the northern Great Basin using predictions from three remotely sensed fractional vegetation cover products: invasive annual grass (IAG) cover from the Sentinel-2-based Rangeland Analysis Platform cover dataset (RAP s2), rangeland annual herbaceous cover (RAH) from the Rangeland Condition Monitoring Assessment and Projection Version 7 (RCMAP v7), and exotic annual grass cover (EAG) from the USGS Exotic Annual Grass product (USGS-EAG).

Inference with respect to the effect of protection was consistent across datasets to the extent that none indicated a significant negative (i.e., suppression of invasion severity) effect of protection within kīpukas. With the exception of RCMAP v7/Set 2, the point estimates for all three datasets instead indicated a positive (i.e., exacerbation of invasion severity) effect of protection, but this effect was accompanied by high uncertainty for all estimates except RAP/Set 1. Broadly, all three fractional vegetation cover datasets support the conclusion that protection from disturbance is ineffective in mitigating the severity of annual grass invasion.

### References

Allred, B. W., B. T. Bestelmeyer, C. S. Boyd, C. Brown, K. W. Davies, M. C. Duniway, L. M. Ellsworth, T. A. Erickson, S. D. Fuhlendorf, T. V. Griffiths, V. Jansen, M. O. Jones, J. Karl, A. Knight, J. D. Maestas, J. J. Maynard, S. E. McCord, D. E. Naugle, H. D. Starns, D. Twidwell, and D. R. Uden. 2021. Improving Landsat predictions of rangeland fractional cover with multitask learning and uncertainty. Methods Ecol. Evol. 12:841–849.

Allred, B. W., S. E. McCord, T. J. Assal, B. T. Bestelmeyer, C. S. Boyd, A. C. Brooks, S. M. Cady, M. C. Duniway, S. D. Fuhlendorf, S. A. Green, G. R. Harrison, E. R. Jensen, E. J. Kachergis, A. C. Knight, C. M. Mattilio, B. A. Mealor, D. E. Naugle, D. O'Leary, P. J. Olsoy, E. S. Peirce, J. R. Reinhardt, R. K. Shriver, J. T. Smith, J. D. Tack, A. M. Tanner, E. P. Tanner, D. Twidwell, N. P. Webb, and S. L. Morford. 2025. Sentinel-2 based estimates of rangeland fractional cover and canopy gap class for the western United States. Sci. Data 12:1889.

Dahal, D., N. J. Pastick, S. P. Boyte, S. Parajuli, M. J. Oimoen, and L. J. Megard. 2022. Multi-species inference of exotic annual and native perennial grasses in rangelands of the western United States using harmonized Landsat and Sentinel-2 data. Remote Sens. 14:807.

Jones, M. O., N. P. Robinson, D. E. Naugle, J. D. Maestas, M. C. Reeves, R. W. Lankston, and B. W. Allred. 2021. Annual and 16-day rangeland production estimates for the western United States. Rangeland Ecol. Manage. 77:112–117.

Rigge, M., C. Homer, L. Cleeves, D. K. Meyer, B. Bunde, H. Shi, G. Xian, S. Schell, and M. Bobo. 2020. Quantifying western U.s. Rangelands as fractional components with multi-resolution remote sensing and in situ data. Remote Sens. (Basel) 12:412.

Rigge, M., C. Homer, H. Shi, D. Meyer, B. Bunde, B. Granneman, K. Postma, P. Danielson, A. Case, and G. Xian. 2021. Rangeland fractional components across the western United States from 1985 to 2018. Remote Sens. (Basel) 13:813.
